# Methodological study of fibroblasts MEF 3T3 heterogeneity cultured on hydrogel substrates with different surface functionalization

**DOI:** 10.1101/2025.06.12.659070

**Authors:** Dawid Wawrzyk, Paweł Goncerz, Zenon Rajfur

## Abstract

Recently, cellular heterogeneity has gained attention in several scientific studies. It’s a phenomenon of the occurrence of differences in cellular parameters, for example, morphology or migration parameters of cells from a single cell line. It has been studied primarily in cancer cells; however, normal cells exhibit analogous behavior. This study investigated morphological and migratory parameters heterogeneity in mouse embryonic fibroblast (MEF 3T3) cells cultured on polyacrylamide (PA) substrates with elasticity of 20 and 40 kPa, which were functionalized with fibronectin solutions in 1, 5, 10 and 20 μg/ml concentrations. Two-hour time lapse microscopy made it possible to quantify the cells spread area and velocity. Based on these parameters and observations of cellular migration strategies, it was possible to classify normal cells into four subpopulations - mesenchymal, amoeboid, slow amoeboid and polygonal/bigonal. Cells grown on softer substrates (20 kPa) showed a greater sensitivity to fibronectin than those grown on stiffer substrates. Cells area was maximal, with the lowest velocity for cells grown on PA substrates functionalized with a 10 μg/ml solution, making their parameters closest to those grown on glass. Cells underwent spontaneous transitions between subpopulations, but no direct transition between mesenchymal and amoeboid or slow amoeboid has been observed. Overall, substrate elasticity and concentration of fibronectin in the functionalizing solution impact the morphology and migration parameters of cells. These findings demonstrate the occurrence of heterogeneity phenomenon in normal mouse fibroblasts. This study also shows the impact of the concentration of fibronectin in the functionalization solution on cells’ behavior.

## Introduction

It is known that significant differences in cell morphology and migration can be noted during the observation of the behavior of theoretically homogenous cell populations^1^. This phenomenon is called cellular heterogeneity. It has primarily been studied in the context of observed behavioral differences occurring in cancer cells of specific cancer types. These differences in subpopulations factors tumor evolution can significantly impact therapeutic efficacy^2,3^. However, the heterogeneity studied in this context has been investigated over longer periods of time, such as days and months. However, this phenomenon was not well studied in terms of short-term changes not directly associated with mutations that may occur over longer periods^4^. Moreover, the effect of heterogeneity in normal cells has been studied to a much lesser extent. As demonstrated, morphological changes of cells, which are not connected with cellular genetics, are also impacted by the physical properties of the microenvironment. For example, the effect of substrate elasticity on which cells are cultured has been well documented^5^. It has also been shown that this factor can influence cellular velocity, area, and shape of migrating cells within subpopulations and dynamics of transitions between subpopulations^4^. This factor also plays a role in a clinical environment, as cancerous tissues exhibit differences in the organization of the extracellular matrix (ECM), contributing to tumor development^6^.

The use of polyacrylamide (PA) substrates allows for the imitation of the environment that cells meet in soft tissues^5^. Its elasticity can be precisely controlled through modifications in the polymer preparation protocol. However, for cells to adhere to the substrate, its surface must be functionalized (coated) with ECM proteins such as fibronectin or type I collagen. It has been observed that the surface density of ECM proteins on polyacrylamide substrates affects the surface and migration speed of cells^7^.

This study focuses on describing how both substrate elasticity and the surface density of fibronectin influence the cellular heterogeneity of mouse embryonic fibroblast cells MEF 3T3. These cells represent a good example of the behavior of non-cancerous adherent cells. The cells’ morphology and migration were measured for 2 hours, allowing the observation of short-term changes in their morphology and migration^4,8^.

## Materials and methods

### Cell culture

MEF 3T3 cells were cultured in DMEM LG medium (Dulbecco’s Modified Eagle’s Medium Low Glucose) (BioWest), with 10% FBS and 1% penicillin and streptavidin, and maintained in a humidified incubator at 37°C and 5% concentration of CO_2_.

### Substrate preparation

Elastic substrates were prepared in glass-bottom Petri dishes (Celvis). Two types of elastic PA substrate with Young’s modulus of 20 and 40 kPa were used. Control cells were grown in glass-bottom Petri dishes without the elastic substrate.

Dishes for PA substrate were first treated with a solution of 3-(trimethoxysilyl) propyl methacrylate (Sigma-Aldrich), acetic acid, and ethyl alcohol (proportion 1:1:14) for 20 minutes at room temperature. Afterwards, dishes were rinsed three times with ethyl alcohol. This process was performed to increase the affinity of PA to glass. PA substrates with a Young modulus of 20 and 40 kPa were made with 40% aqueous solution of acrylamide (Bio-Rad), 2% aqueous solution of bis-acrylamide (Bio-Rad), and 10 mM aqueous solution of HEPES (Sigma-Aldrich). Proportions of acrylamide, bis-acrylamide and HEPES were selected according to the standard procedure^9^. To start the process of polymerization, 10% ammonium persulfate (Bio-Rad) and tetramethylethylenediamine (BioShop) in ratios of 1/200 and 1/2000 to the total volume of the mixture were added. 22 μl of the mixture was put on each dish and covered with a coverslip of 18 mm diameter. After an hour, substrates were covered with PBS, coverslips were removed, and dishes were put in a fridge for 24 hours. Then, the substrate was activated with SulfoSANPAH solution (Thermo Scientific) for 5 minutes under ultraviolet light. Afterwards, the substrate was washed 3 times with 10 mM aqueous solution of HEPES and 3 times with sterile PBS. In the last step, substrates were incubated for 24h at 4°C, respectively, with a 1, 5, 10, and 20 μg/ml fibronectin solution. Lastly, the substrate was washed with PBS.

### Observation of cell migration

Cell migration and morphology were recorded using an inverted Axio Observer.Z1 microscope (Zeiss), AxioCam camera (Zeiss), and a Plan Apochromat 10×/0.45 objective (Zeiss). Time-lapse measurements were carried out with the Definite Focus component (Zeiss) used to maintain a proper focal plane. Images were acquired for 2 hours with 5-minute interval between consecutive frames. Recording started 1 hour after cell seeding, cells were kept at a temperature of 37°C and 5% concentration of CO_2_.

### Image analysis

From every substrate, 48 cells were chosen to analyze. To reduce the amount of data, every other frame was used for the analysis of cell area and velocity. However, all frames were analyzed to assess the type of cell migration

Cell morphology was analyzed manually using the ROI Tracker plugin^10^ in the ImageJ software and subsequently converted to binary masks. The binary masks were then analyzed using ImageJ software to calculate the area and centroid location of every cell. Cell velocity was defined as the change in the centroid position between two consecutive frames.

For a cell to be selected for analysis, it had to meet the following conditions during all times of registration: (1) it was clearly visible in the picture, (2) it was not undergoing cell division, (3) it was isolated (not being in contact) from other cells.

### Subpopulations classification

Cells were classified into four subpopulations based on migration parameters, their shape and its change, and their area. Baseline in defining subpopulations was defined for Walker WC256 cells^4^ and adjusted for better representation of MEF 3T3 cells.

Cells qualified as bigonal or polygonal were stretched in two (bigonal) or more (polygonal) directions and poorly spread (Fig 1). Their shape was stable and changed slowly by extending lamellipodia. This process was the only one that resulted in the change of centroid position.

**Figure 1.**
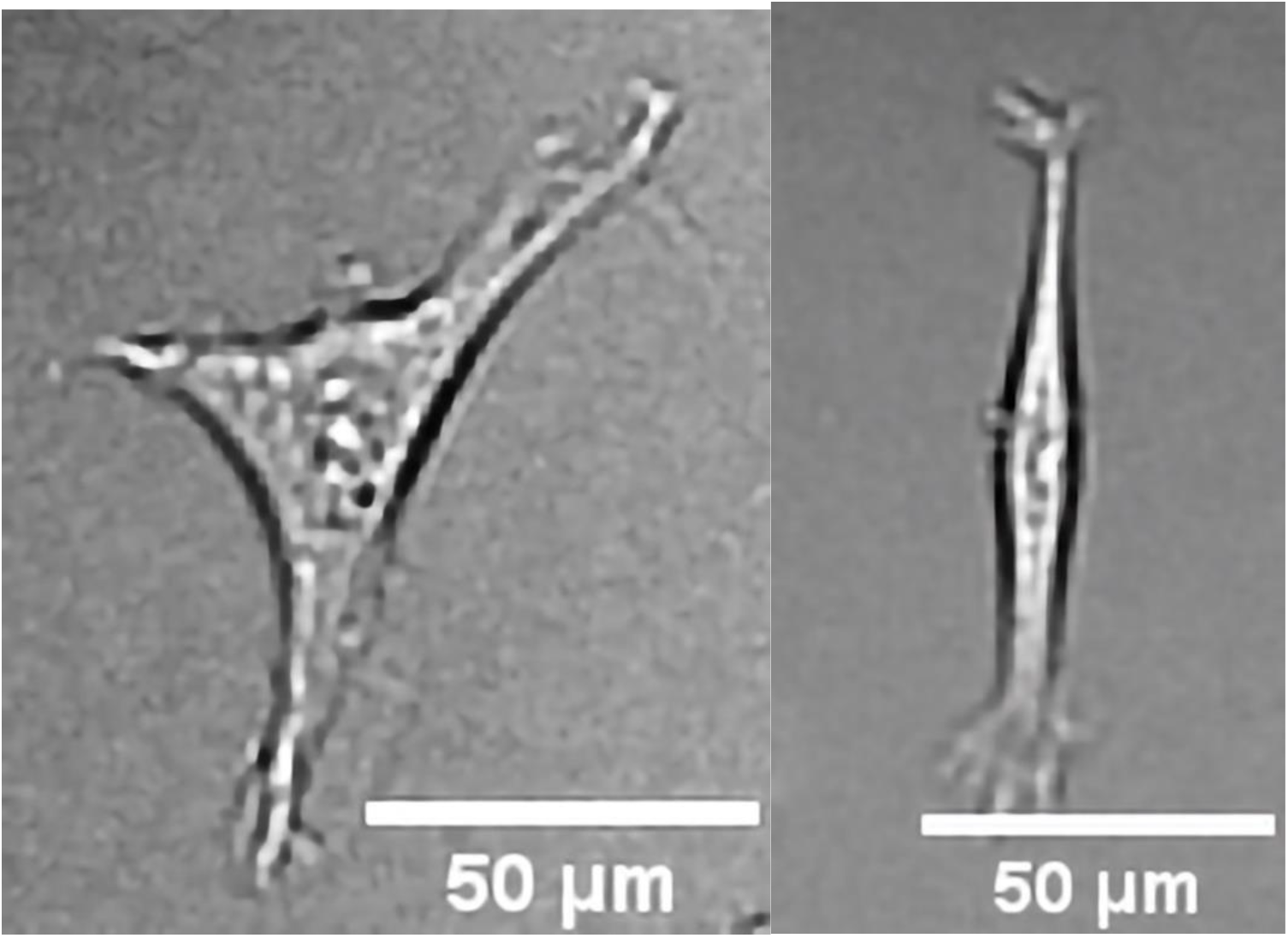
Examples of polygonal (left panel) or bigonal (right panel) cells.

Cells qualified as mesenchymal were widely spread (Fig. 2). Two modes of cell migration were observed. The first mode was realized by spreading and stretching of the leading edge coordinated with a detachment of the opposite edge, resulting in a relatively stable shape. The second, a slower type, was a keratinocyte-like gliding with no extended leading edge.

**Figure 2.**
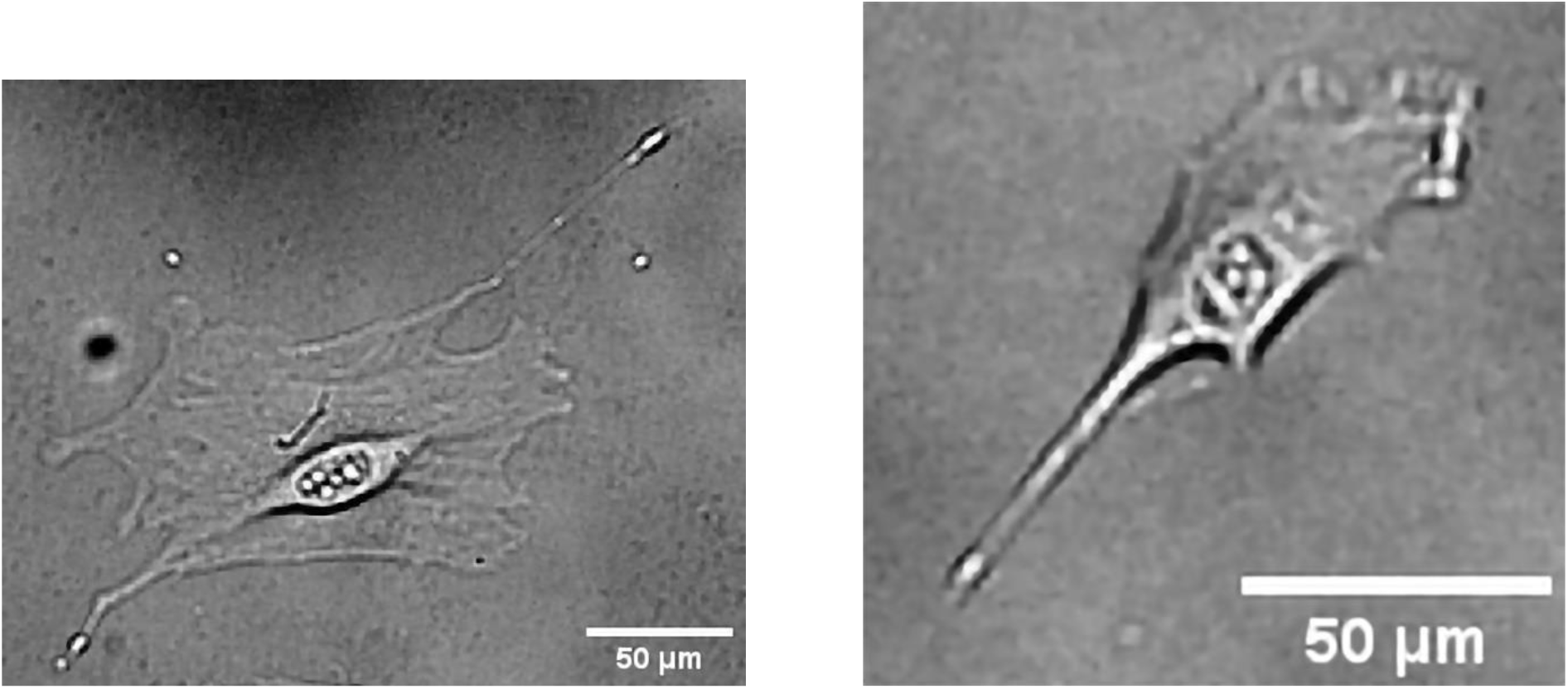
Examples of mesenchymal cells.

Cells qualified as amoeboid exhibited quickly changing shape, with a relatively small area (Fig. 3). Their migration resulted from cycles of spreading of one edge before detachment of the majority of the cell, resulting in a fast change of shape and centroid position. Alternatively, the movement was caused by the formation of a round protrusion on the leading edge, leading to a fast change of position of the centroid.

**Figure 3.**
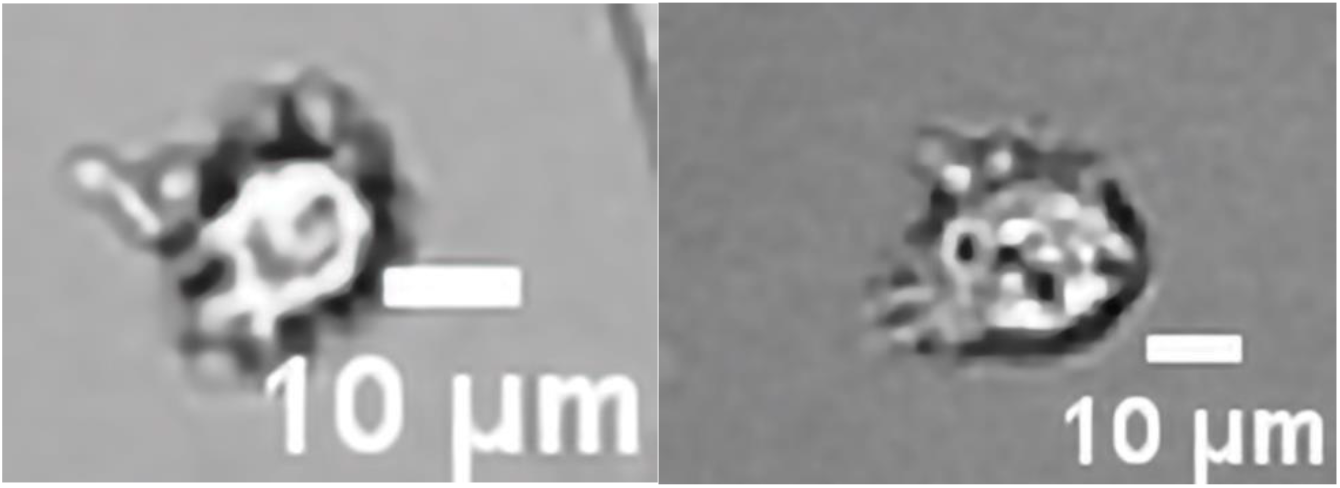
Examples of amoeboid cells.

One more subpopulation was observed. Cells qualified as slow amoeboid were the least spread with a round shape like the amoeboid with quickly forming and disappearing protrusions, which did not lead to a significant change in the centroid position.

### Statistical analysis

Further analysis was performed using Microsoft Excel and OriginPro 2021. To describe data, median and interquartile ranges (IQRs) were used because of large differences in parameters between cells in every group. Mann–Whitney test was used to check statistical differences between data series.

## Results

### Influence of fibronectin concentration in the functionalizing solution of a cellular substrate on migration and morphology parameters of MEF 3T3 cells

This study examined the effect of functionalization of substrates surface with different concentrations of fibronectin. To investigate possible differences between cells cultivated on substrates with different elasticity (Young modulus) and functionalized with different fibronectin concentration, the focus was on two parameters: cellular area and velocity. At first, the entire cell population was considered without considering their heterogeneity (Fig. 4).

**Figure 4.**
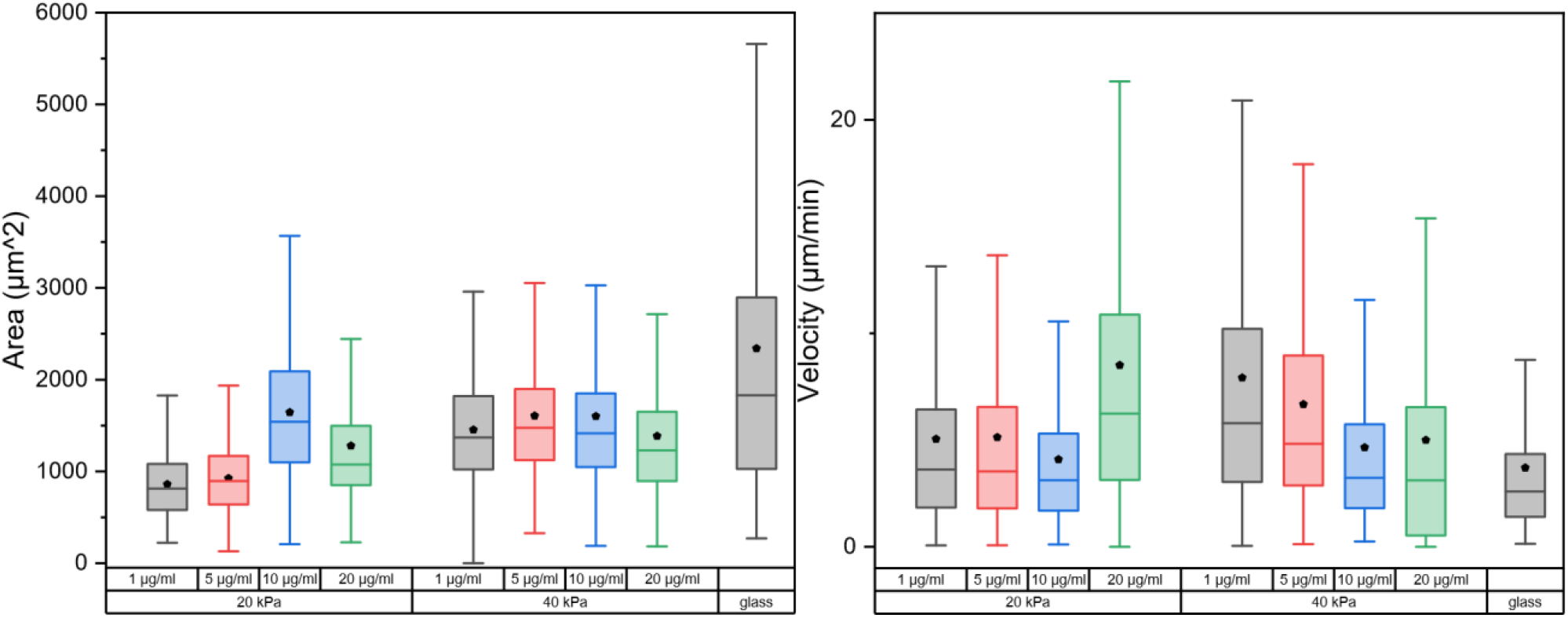
The influence of fibronectin concentration in functionalizing solution on a cellular substrate with Young modulus of 20 (left panel) and 40 kPa (right panel) on the area and velocity of MEF 3T3 cells.

Fibronectin concentration in the functionalization solution had a bigger impact on cells cultured on softer substrates. Cells grown on a substrate with Young’s modulus of 20 kPa functionalized with 10 μg/ml fibronectin solution were the most spread (Fig. 4). Changing the fibronectin concentration from 10 mg/ml in the solution led to a decrease in cell surface area.

Stiffer substrates (40 kPa) exhibited a similar relationship. However, the differences were much smaller, statistically significant only for the decrease in surface area for concentrations 20 and 1 μg/ml. Maximal area was observed on substrates functionalized with 5 and 10 μg/ml fibronectin solution (Fig. 4).

Velocity showed the inverse relationships, with the lowest value for 10 μg/ml fibronectin solution, for both stiffnesses.

On both substrates functionalized with 10 μg/ml fibronectin solution combination of parameters (cellular area and velocity) was closest to the control cells grown on glass. Data analyzed from a whole cell population showed large dispersion in both parameters values.

### Morphological heterogeneity of MEF 3T3 cells

Cells were categorized based on morphology and migration strategies as shown in the “Materials and methods” section. Our observations showed that there is a portion of cells with morphological similarity to the amoeboid subpopulation, but they did not exhibit rapid displacement. Because of that, they were categorized as “slow amoeboid<u>”</u>. We hypothesize that the differences in migration are not caused by changes within the cells, but by local parameters of substrate, such as ECM protein concentration or surface topography, that impacted the formation of points of adhesion.

Additionally, the analysis showed that on glass and substrates with a Young’s modulus of 40 kPa functionalized with 5 or 10 μg/ml fibronectin, as well as on substrates with 20 kPa stiffness functionalized with 10 μg/ml fibronectin, cells appeared more spread and morphologically resembled mesenchymal subpopulations; however, they exhibited significantly lower velocity. Due to meeting the criteria, they were classified as mesenchymal.

### Distribution of MEF 3T3 cells subpopulations and cellular transitions between them

To determine the distribution of cells into subpopulations, we calculated percentage of cells of each subpopulation on all frames (Fig. 5). For every substrate majority of cells were classified as mesenchymal except for 20 kPa Young modulus functionalized with 5 μg/ml fibronectin solution where mesenchymal and polygonal/bigonal subpopulations appeared to be equal. Subpopulation distribution of cells grown on elastic substrates functionalized with 10 μg/ml fibronectin solution shows close resemblance to cells grown on glass. Changes in fibronectin concentration clearly impacted subpopulations distribution on elastic substrates by reducing the share of mesenchymal subpopulation in favor of other subpopulations when the concentration of fibronectin differed from 10 μg/ml. This effect was less pronounced on stiffer substrates, where only the substrate functionalized with 20 μg/ml of fibronectin solution clearly differed from others.

**Figure 5.**
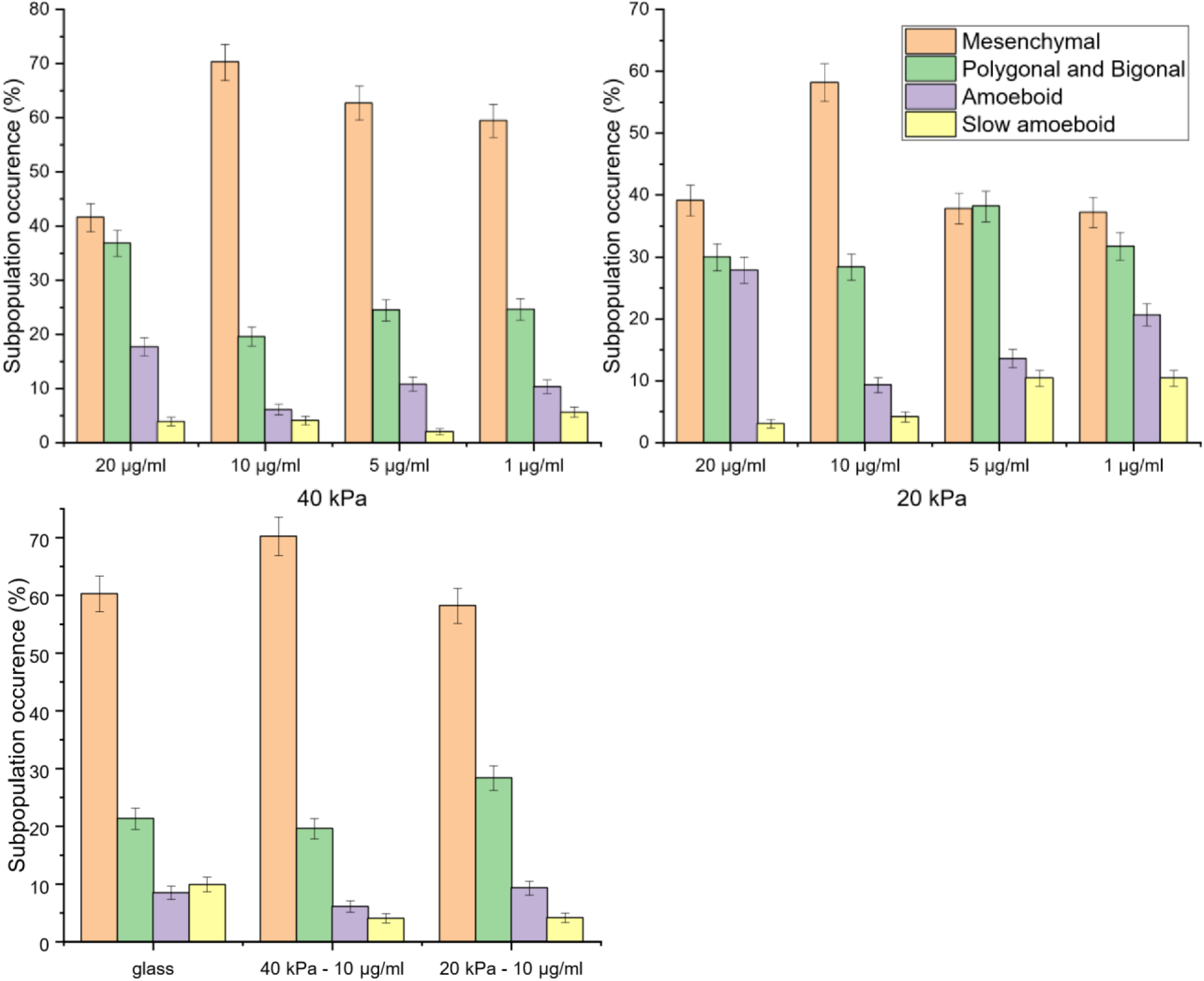
The occurrence of different subpopulations of MEF 3T3 cells on substrates of different elasticity and functionalized with solutions with different concentration of fibronectin.

Not every cell belongs to only one population during the observation time (2h), transitions between subpopulations occurred without external stimulation. Interestingly, some cells could revert to their initial subpopulation after transition. For example, cells initially classified as mesenchymal have been observed to transition to polygonal/bigonal and then to an amoeboid or slow amoeboid phenotype and vice versa. What is important, no direct transition from mesenchymal to amoeboid (or vice versa) was observed (Fig. 6).

**Figure 6.**
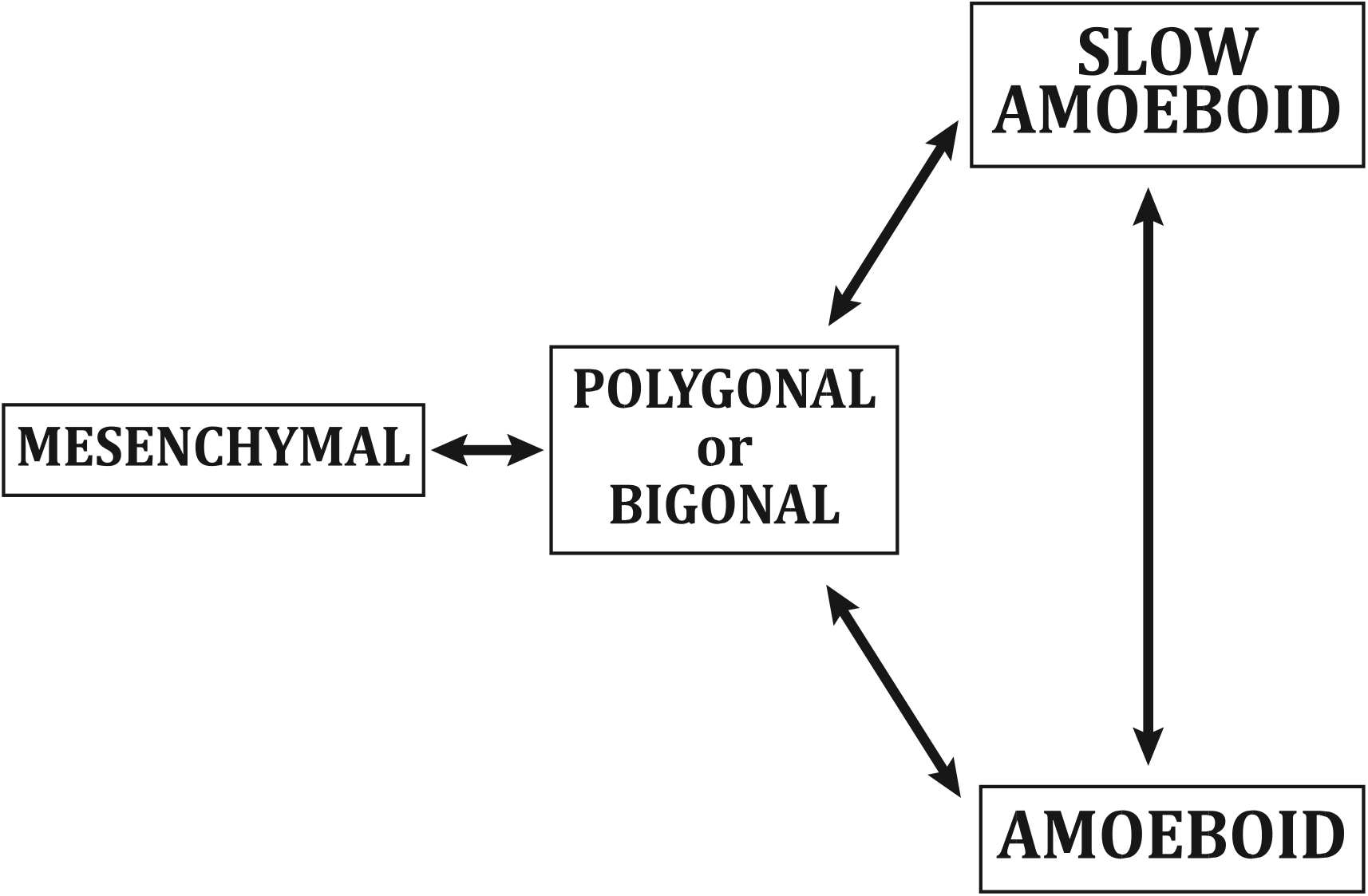
Diagram of observed transitions between subpopulations.

The majority of transitions for all investigated substrates occurred between mesenchymal and bigonal/polygonal and between amoeboid and bigonal/polygonal. Transitions that included a slow amoeboid state were less common (Figure 7). Most cells never went through transitions between subpopulations during the observation time.

**Figure 7.**
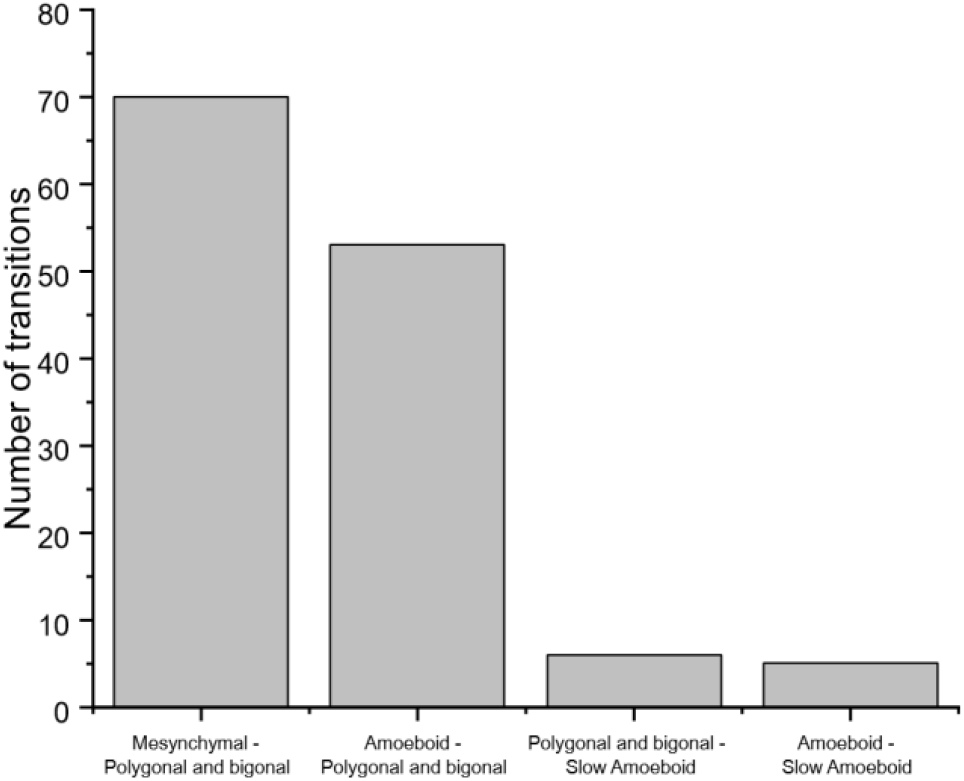
Total number of transitions between subpopulations on every substrate.

### Influence of cellular substrate elasticity and surface functionalization on morphology and migration of MEF 3T3 subpopulation

Subsequent analysis of the previously described parameters, taking into consideration cellular division into subpopulations, was carried out. It allowed us to determine whether the impact on velocity and area is only due to the change in the proportion of subpopulations or is independent of it. The parameters describing the most numerous mesenchymal subpopulation are close to the whole population (Figure 8 and Figure 4). These results show the influence of elasticity on mesenchymal population and that fibronectin concentration has no real influence on the area of cells grown on 40kPa substrate.

**Figure 8.**
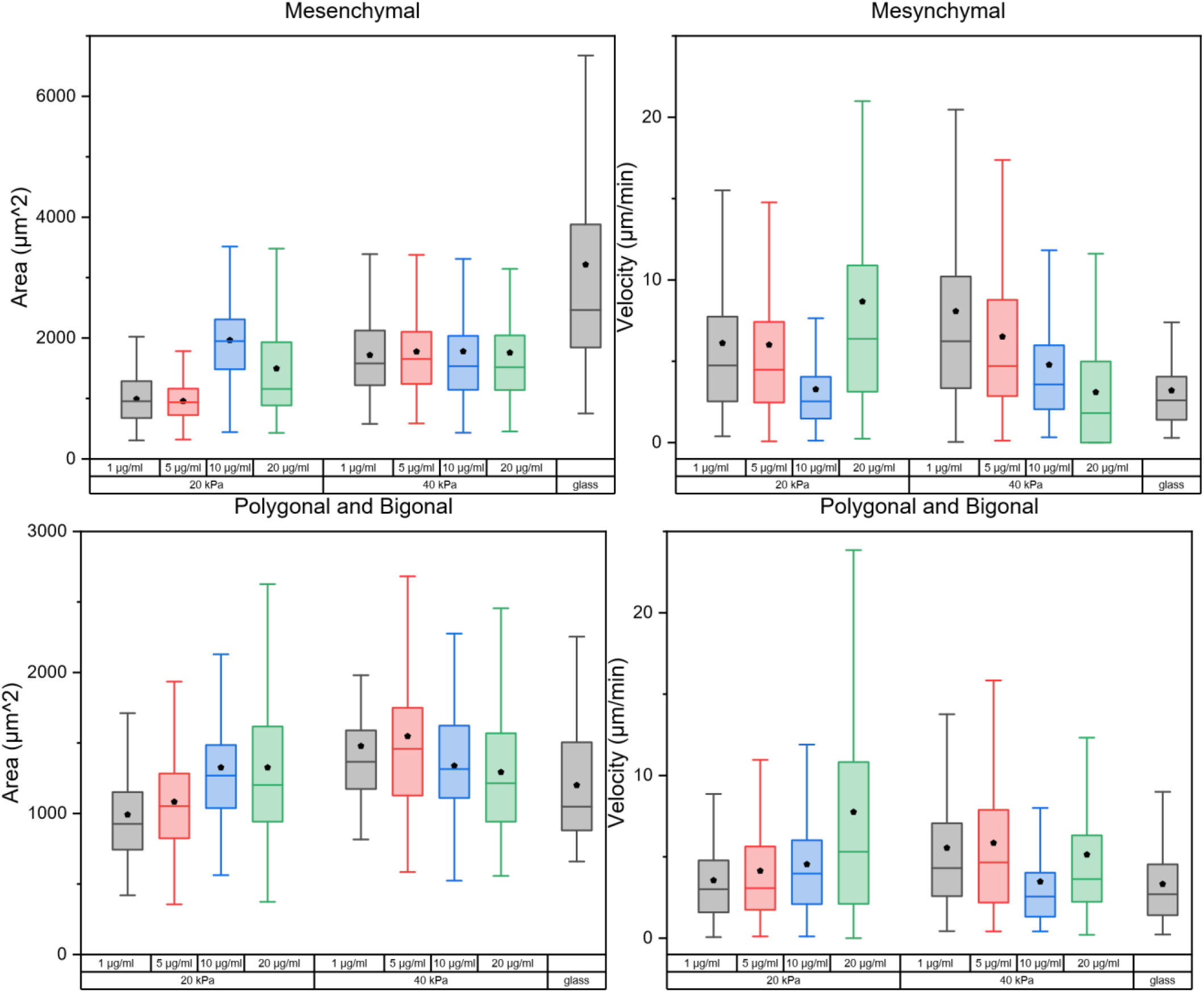
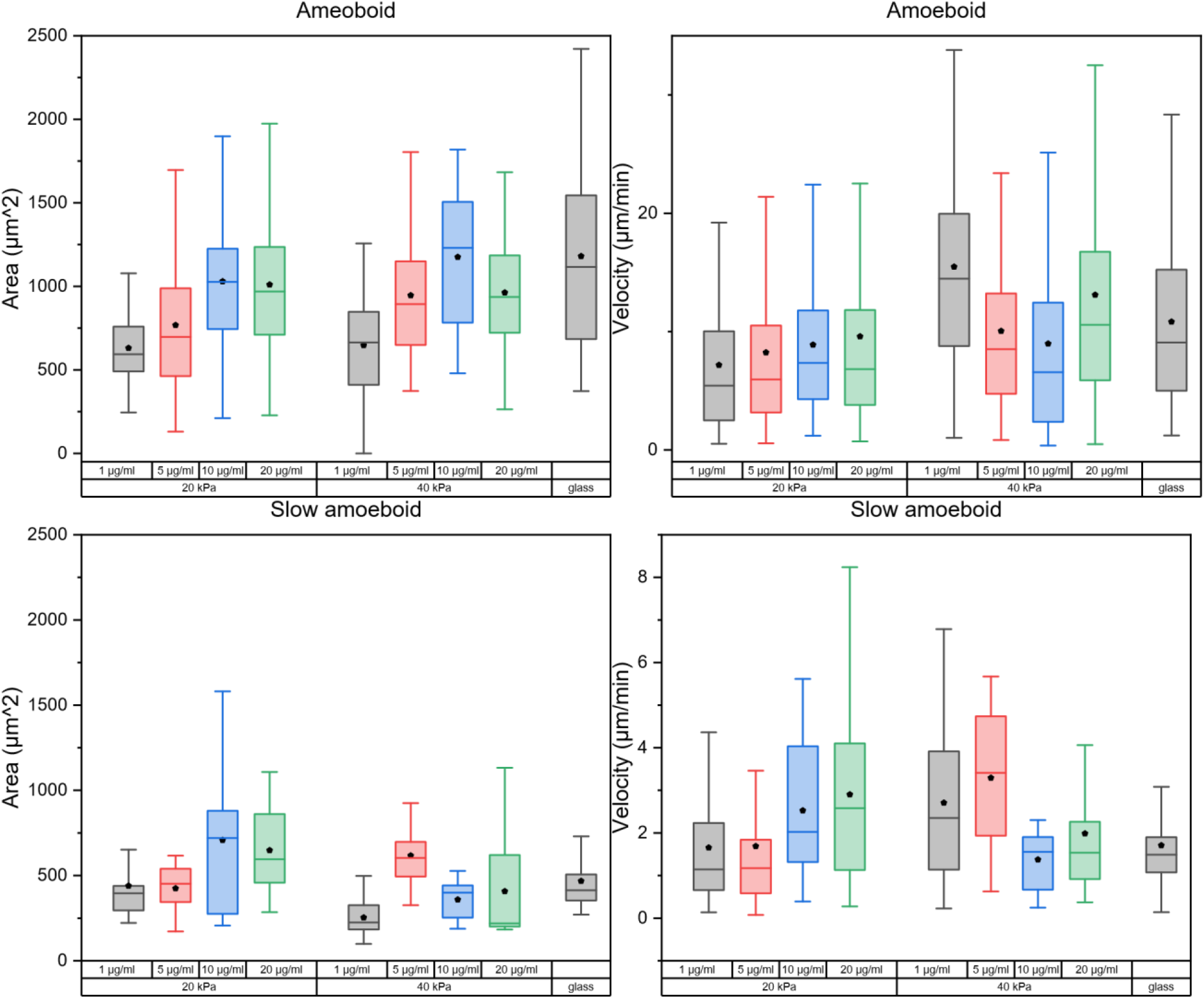
The influence of fibronectin concentration in functionalizing solution on a cellular substrate with Young’s modulus of 20 and 40 kPa on area and velocity of mesenchymal(a), polygonal and bigonal(b), amoeboid(c), and slow amoeboid (d) of MEF 3T3 cell line.

However, the migration velocity is influenced by fibronectin concentration (Figure 8a). For polygonal and bigonal subpopulations, changes were similar to the mesenchymal subpopulation (Figure 8b). The cell’s area of the amoeboid subpopulation is influenced by fibronectin concentration, even on a 40 kPa substrate, while velocity is much less impacted. For the slow amoeboid subpopulation it is not clear what is the impact of substrate elasticity or its functionalization on the cellular parameters. For all subpopulation<u>s</u> (excluding slow amoeboid) substrate functionalized with 10 μg/ml fibronectin solution, their parameters were the closest to cells cultured on glass.

## Discussion

The current study showed that the normal cell line (MEF 3T3) is not homogenous but exhibits the presence of specific subpopulations. Therefore, it is necessary to take into consideration that cells can belong to different subpopulations to make analysis of their behavior more precise. In this study, the parameters of cellular area and velocity of MEF 3T3 were characterized, taking into consideration cell line heterogeneity. It was shown that the morphology and migration strategy of MEF 3T3 cells are interrelated, and partition of cells into subpopulations can occur during measurement. Subpopulations could be defined as described in ^4^ without significant changes. It can be speculated that similar categorization could also be used in the case of other adherent cell lines. Similarly to cancer cells, the mesenchymal subpopulation was most numerous on glass, but changes in distribution in population were more connected to fibronectin concentration in the substrate functionalizing solution. Therefore, significant changes shown in previous studies^4^ could differ on substrates functionalized with different fibronectin concentrations.

The current study found that area, velocity and occurrence of subpopulation<u>s</u> of MEF 3T3 cells are impacted by fibronectin concentration in the functionalizing solution of a cellular substrate. It has been shown that differences in substrate coating with ECM proteins may lead to different results, especially with less stiff substrates, where the impact on area, velocity, and distribution into subpopulations was greater. Cells cultured on PA substrates functionalized with 10 μg/ml of fibronectin solution show the closest resemblance to the cells cultured on glass, including the highest area, the lowest velocity, and a high percentage of mesenchymal subpopulation, followed by bigonal and polygonal subpopulations.

Observed differences in cell velocity and spread area may be caused by the differences in the number of places where the cell can adhere. If the density of fibronectin on the substrate surface is high, available integrin receptors could reach the limit on the lower surface. When the density of fibronectin on the surface is too low, cells cannot spread effectively due to limited availability of adhesive sites^7^. Differences in velocity could have the same source. For amoeboid cells, migration requires the creation of a smaller number of adhesion points than for other subpopulations, therefore, they are less impacted by the density of available adhesion sites^11^. The velocity of mesenchymal and polygonal/bigonal subpopulations is similar, but its source is different. Change of centroid positions in mesenchymal subpopulation is caused by migration of a whole cell, while in polygonal and bigonal subpopulations it is caused by stretching and retraction of lamellipodia.

Transitions between subpopulations occurred spontaneously and were reversible. Most common transitions occurred between mesenchymal and polygonal/bigonal and between amoeboid and polygonal/bigonal subpopulations. No transitions between amoeboid or slow amoeboid and mesenchymal subpopulations were observed. Transitions like these in other studies were triggered with external chemical or physical stimuli in cancer cells^11,12^.

## Conclusions

Our study showed that normal cell line shows the occurrence of heterogeneity effect in a way similar to the cancerous cell line^4^. Not only stiffness but also the concentration of fibronectin solution used to coat the substrate surface have an impact on that phenomenon.

The current study involved a 2 h observation with time-lapse technique with sampling every 10 minutes. Sampling time has the biggest impact on amoeboid subpopulation, because it contains cells with fast-changing direction of migration. Inclusion of turning angle in migration description of cells^13^, ratio of area to perimeter, and speed of surface changes could also lead to a more accurate description.

## Author Contributions

DW - Investigation, Methodology, Formal Analysis, Visualization, Writing – Original Draft ; PG - Formal Analysis, Writing – Review & Editing; ZR - Conceptualization, Methodology, Project Administration, Supervision, Writing – Review & Editing

## References

1 Welch DR. Tumor heterogeneity—a ‘contemporary concept’ founded on historical insights and predictions. Cancer Res. 2016;76(1):4–6.

2 Jacquemin V, Wilson CR, McGhee E, et al. Dynamic cancer cell heterogeneity: Diagnostic and therapeutic implications. Cancers. 2022;14(2):280.

3 Dagogo-Jack, Ibiayi, and Alice T. Shaw. “Tumour heterogeneity and resistance to cancer therapies.” Nature reviews Clinical oncology 15.2 (2018): 81–94.

4 Mielnicka A, Strąkowska M, Nowak A, et al. Impact of elastic substrate on the dynamic heterogeneity of WC256 Walker carcinosarcoma cells. Sci Rep. 2023;13(1):15743.

5 Pelham Jr, Robert J., and Yu-li Wang. “Cell locomotion and focal adhesions are regulated by substrate flexibility.” Proceedings of the national academy of sciences 94.25 (1997). 13661–13665.

6 Marusyk A, Janiszewska M, Polyak K. Intratumor heterogeneity: the rosetta stone of therapy resistance. Cancer Cell. 2020;37(4):471–484.

7 Gaudet C, Ray SL, Drake B, et al. Influence of type I collagen surface density on fibroblast spreading, motility, and contractility. Biophys J. 2003;85(5):3329–3335.

8 Bergert M, Zieseniss A, Eisenhardt A, et al. Cell mechanics control rapid transitions between blebs and lamellipodia during migration. Proc Natl Acad Sci U S A. 2012;109(36):14434–14439.

9 Tse JR, Engler AJ. Preparation of hydrogel substrates with tunable mechanical properties. Curr Protoc Cell Biol. 2010;47(1):10.

10 Entenberg D, Condeelis J. ROI Tracker [software]. Supplied by David Entenberg and John Condeelis; supported by CA100324 and GM064346.

11 Brábek J, John CM, Kára DA, et al. The role of the tissue microenvironment in the regulation of cancer cell motility and invasion. Cell Commun Signal. 2010;8:1.

12 Carragher NO, Frame MC, Fincham VJ, et al. Calpain 2 and Src dependence distinguishes mesenchymal and amoeboid modes of tumour cell invasion: a link to integrin function. Oncogene. 2006;25(42):5726– 5740.

13 Kołodziej T, Żurada M, Lewandowski M, et al. Morphomigrational description as a new approach connecting cell’s migration with its morphology. Sci Rep. 2023;13(1):11006.

